# sAMP-VGG16: Drude polarizable force-field assisted image-based deep neural network prediction model for short antimicrobial peptides

**DOI:** 10.1101/2023.06.04.543607

**Authors:** Poonam Pandey, Anand Srivastava

## Abstract

During the last three decades, antimicrobial peptides (AMPs) have emerged as a promising therapeutic alternative to antibiotics. The approaches for designing AMPs span from experimental trial-and-error methods to synthetic hybrid peptide libraries. To overcome the exceedingly expensive and time-consuming process of designing effective AMPs, many computational and machine-learning tools for AMP prediction have been recently developed. In general, to encode the peptide sequences, featurization in these rely on approaches based on (a) amino acid composition, (b) physicochemical properties, (c) sequence similarity, and (d) structural properties. In this work, we present an image-based deep neural network model to predict AMPs, where we are using feature encoding based on Drude polarizable force-field atom types, which can capture the peptide properties more efficiently compared to conventional feature vectors. The proposed prediction model identifies AMPs with promising accuracy and efficiency and can be used as a next-generation screening method for predicting new AMPs. The source code is publicly available at the Figshare server sAMP-VGG16.

## I. INTRODUCTION

Antimicrobial peptides (AMPs) constitute a diverse repertoire of antibacterial agents and therefore a valuable resource for drug discovery [1–3]. During the last three decades, AMPs have emerged as a promising therapeutic alternative to antibiotics due to their unique mechanism of action to outwit bacterial resistance. Particularly, cationic short AMPs (sAMPs) have sparked great interest in recent years for their potential in the development of novel antibacterial drugs. Unlike traditional antibiotics, sAMPs eliminate or inhibit bacteria mainly via a membrane-active mechanism, which does not require either site-specific attachment or interference in bacterial metabolism [4]. Despite their potential, there are several hurdles in using sAMPs as pharmaceutical agents. sAMPs often have poor stability, specifically their proteolytic stability; may have high-level of toxicity, and can also have high cost required for production [5]. While it is possible to overcome the first two limitations by rigorous optimization and peptide engineering, the high cost of manufacturing renders it prohibitively expensive to screen a large number of peptides [6]. Therefore, the cost of peptide screening plus the potential need to improve the stability and toxicity of sAMPs make it unappealing for research labs and pharmaceutical corporations.

In the last decades, multiple efforts have been made to collect databases of AMPs and to classify these protein sequences through various computational techniques [7–14]. Notable recent advances in computing power and statistical learning tools have rendered supervised machine learning a promising strategy for harnessing large datasets for the high-throughput screening of antimicrobial peptides (AMPs). Here we provide a brief chronological summary of the developements made in the field over the last 10-15 years. In 2007, Lata et al. proposed an AMP classification tool based on the quantitative matrix (QM), artificial neural network (ANN), and support vector machine (SVM) [15]. Later on, Jenssen et al. developed a mathematical model for AMP prediction using contact energy descriptors for each pair of amino acids, 78 biophysical inductive and conventional QSAR descriptors [16, 17]. In 2008, Fjell et al. developed a profile hidden Markov model (HMM) based AMPer resource to recognize AMPs [18]. Later on, in 2009, Chersakov et al. utilized a high-throughput screening-assisted Artificial Neural Network (ANN) model to discover potent AMPs effective against multi-drug resistant bacteria[19]. In 2010, Thomas et al. developed CMAP database along with ML-based AMP prediction tools using random forest (RF), SVM, and discriminant analysis (DA) [20]. In 2011, Wang et al. developed a more efficient web server by integrating the BLASTP-based sequence alignment method and the feature selection method with amino acid composition (AAC) and pseudo amino acid composition (PseAAC) [21]. In the same year, Torrent et al. developed an ANN-based model using physicochemical properties derived from peptide sequences to predict AMPs and to assess their antimicrobial potency [22]. In 2013, Maccari et al. proposed a RF-based model to design and validate the AMPs with both natural and non-natural amino acids[23]. In the same year, Xio et al. proposed a fuzzy K-nearest neighbor (FKNN) based method using pseudo amino acid composition [24]. In 2015, Giguere et al. developed a graph-based algorithm to predict AMPs for ML predictors using generic string (GS) kernel [25]. In 2016, Schneider et al. proposed self-organizing maps (SOM) based deep network to analyze AMPs[26]. In 2017, Meher et al. developed a more efficient SVM-based AMP prediction model using PseAAC feature vector by incorporating compositional, physicochemical and structural features[27]. In 2018, Bhadra et al. developed the AmPeP model using RF and distribution patterns of amino acid properties[28]. Vishnepolsky et al. proposed a semi-supervised ML approach utilizing the DBSCAN algorithm and physicochemical descriptors[29]. In the same year, Veltri et al. proposed a neural network model with convolutional and recurrent layers by utilizing primary sequence information[30]. In 2020, Yan et al. developed a convolutional neural network (CNN) and RF-based prediction model for short-antimicrobial peptides (≤ 30 AA)[31]. In 2022, Yan et al. developed sAMPpred-GAT, a graph attention network (GAT) based on peptide sequence, structure, and evolutionary information[32]. For sake of brevity, we have also summarized the above review in the form of a table (see Table 1).

**TABLE I:**
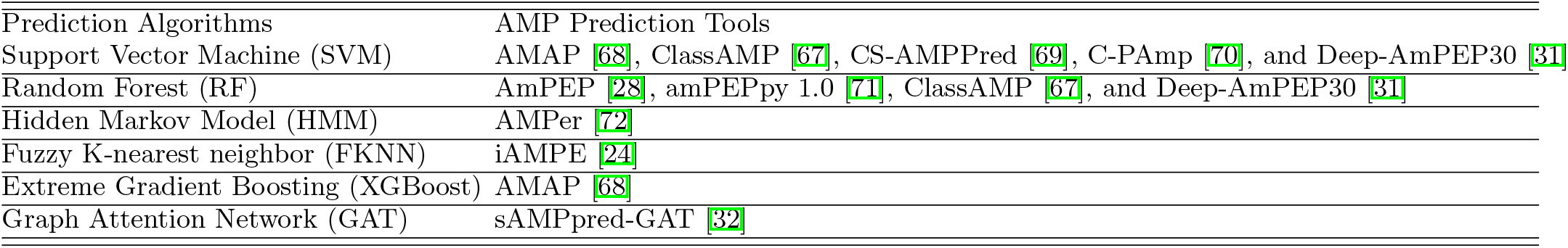
Details of prediction algorithms used for AMP prediction.

| Prediction Algorithms | AMP Prediction Tools |
| --- | --- |
| Support Vector Machine (SVM) | AMAP [68], ClassAMP [67], CS-AMPPred [69], C-PAMP [70], and Deep-AmPEP30 [31] |
| Random Forest (RF) | AmPEP [28], amPEPpy 1.0 [71], ClassAMP [67], and Deep-AmPEP30 [31] |
| Hidden Markov Model (HMM) | AMPer [72] |
| Fuzzy K-nearest neighbor (FKNN) | iAMPE [24] |
| Extreme Gradient Boosting (XGBoost) | AMAP [68] |
| Graph Attention Network (GAT) | sAMPpred-GAT [32] |

**TABLE II:**
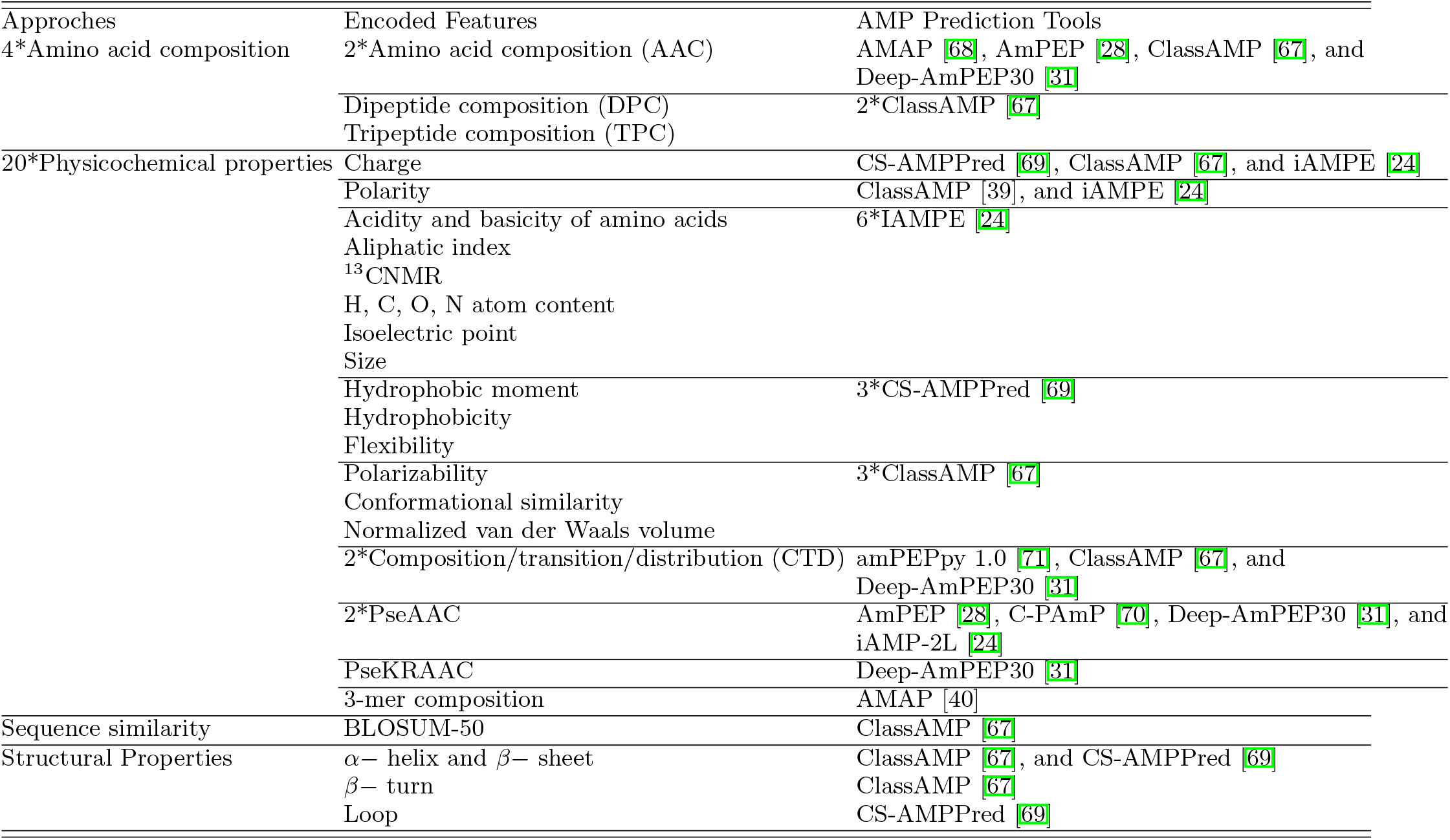
Details of peptide/protein encoding approaches used for AMP prediction

In light of recent advances in the discovery of sAMPs ((≤ 30 AA), exhibiting potent antibacterial properties, high specificity, and low toxicity has prompted an exciting major shift in the field of AMP research[33–35]. Notable the shorter length of these AMPs renders synthesis, modification, and optimization straightforward and more affordable than with conventional AMPs. The rapid advancement facilitated by the usage of sAMPs has made them an intriguing and cost-effective class of biomolecules for drug screening[35]. The computational approaches consist of two fundamental steps: [i] Feature Extraction; and [ii] prediction model development and evaluation. In the feature extraction phase, several discriminative features are proposed to encode protein/peptide sequences. In prior studies, to encode the protein sequences, featurization were depends on the approaches based on (a) amino acid composition, (b) physicochemical properties, (c) sequence similarity, and structural properties. The prediction models were constructed using traditional ML algorithms such as Support Vector Machine (SVM), Random Forest (RF), decision tree (DT), Deep learning, ensemble learning methods, and Graph Attention Network (GAT) method. The details of peptide/protein encoding methods are given in Table II.

MD simulation studies using various force fields play a key role to comprehend chemical and biological systems. The effectiveness of the MD simulation studies in terms of the accuracy of the anticipated attributes is influenced by the simulation algorithms, the complexity of the systems, and most significantly the underlying force field. The Drude polarizable FF[36–38] is a well-known atomistic FF is well-established atomistic FF that incorporates classical Drude oscillators to accommodate electronic polarization. The Drude polarizable FF parameters have been meticulously optimized to model biological systems, such as proteins[39–42], lipid[43–45], nucleic acids[46–48], carbohydrates[49–52], ions[45, 53], and small molecules[54–56] incorporated in the systems. In this study, the Drude atom type and their connectivity information were used to generate input features for sAMP prediction model. In Drude FF, an atom can adopt a distinct atom-type definition premised on its connectivity along with the formal charge state. Atom-type information has an advantage over the usage of amino acidresidue-based feature vectors because it differentiates atoms depending on their chemical environment in the peptide residues. Therefore, the usage of atom-type information in feature extraction will inherently encompass both local and non-local effects, providing effective atomistic features for the prediction model. This paper proposed an efficient novel sAMP-prediction model, based on atomistic feature information. In this work, Deep-AmPEP30 training and validation dataset has been used for 1 to 1 comparison to develop a more precise and robust prediction model for sAMPs.

To the best of our knowledge, this is the first time that FF-based atomistic information in conjunction with VGG16 based DNN model is proposed for the protein classification problem. The previously reported ML algorithms were based on amino acid residue-based feature vectors; like AA composition, physicochemical properties, etc.; combined with traditional classifiers like SVM, RF, DT, deep learning, and ensemble learning methods. Consequently, the proposed prediction model is of great interest and appears to be very promising, not only because of the application of atomistic information but also because of the transfer-learning methods, which require less tuning of the hyperparameters to build effective prediction models for peptide classification.

## II. METHODOLOGY

The present study intended to predict short amino acid peptides (sAMPs), defined as peptides with a sequence length of amino acids less than or equal to 30 amino acids. The protocol for developing the VGG16based DNN model to predict the AMPs using Drude FF atom types is divided into three parts: [1] Dataset preparation, [2] Feature extraction and image generation for VGG16-AMP model, [3] Training and validation of VGG16-sAMP model.

### A. Dataset Description

The training and validation dataset were collected from the previously reported study by Yan et al.31 The training dataset consists of 1529 sAMPs and 1529 non-sAMPs. Therefore, the final balanced training dataset comprises 3058 peptide sequences. A benchmark dataset from Yan et al. study was used to evaluate the proposed prediction model, as well as to compare the proposed prediction model to existing state-of-the-art prediction models. The benchmark dataset consists of 94 sAMPs and 94 non-sAMPs. The dataset was processed by CD-HIT for the sake of redundancy removal at a threshold of 0.8, i.e., the sequence which showed similarity of more than 90 % were excluded. As this preprocessing was already performed, no such step was repeated in this study. The training and benchmark dataset information is provided in Figure 1.

**FIG. 1:**
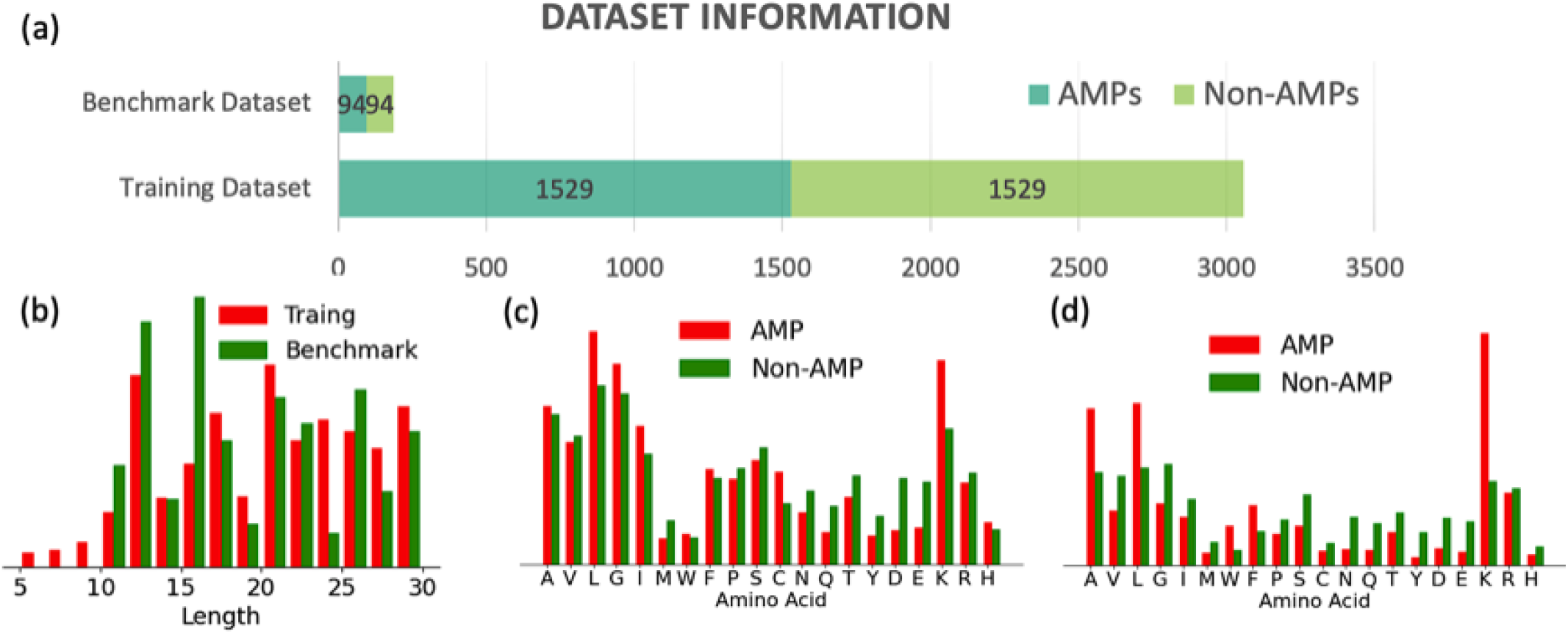
Overview of the training and benchmark dataset: (a) sAMP and non-sAMP distribution in training and benchmark dataset; (b) Comparison of the peptide length distribution in training and benchmark dataset; (c) Comparison of the amino acid distribution in sAMPs and non-sAMPs in training dataset; (d) Comparison of the amino acid distribution in sAMPs and non-sAMPs in benchmark dataset

### B. Feature Extraction and Image Generation for VGG16-AMP Model

In this work, feature encoding is based on Drude polarizable FF atom types. In the first step, psf file has been generated for each input peptide sequence using protein Drude polarizable FF in CHARMM. In the second step, the psf file is used to extract feature information about atom type connectivity, which includes bond [1-2], angle [1-3], and dihedral [1-4] connectivity. In the third step, the three connectivity matrices are normalized in a range of 0-255 to generate three actual uniform channels. In the fourth step, the three channels were merged to create a 3channel image for each peptide sequence. In this way, all sequences are transformed into 3-channel images, ready to forward to target convolutional neural networks. The framework of feature extraction and image generation is provided in Figure 2.

**FIG. 2:**
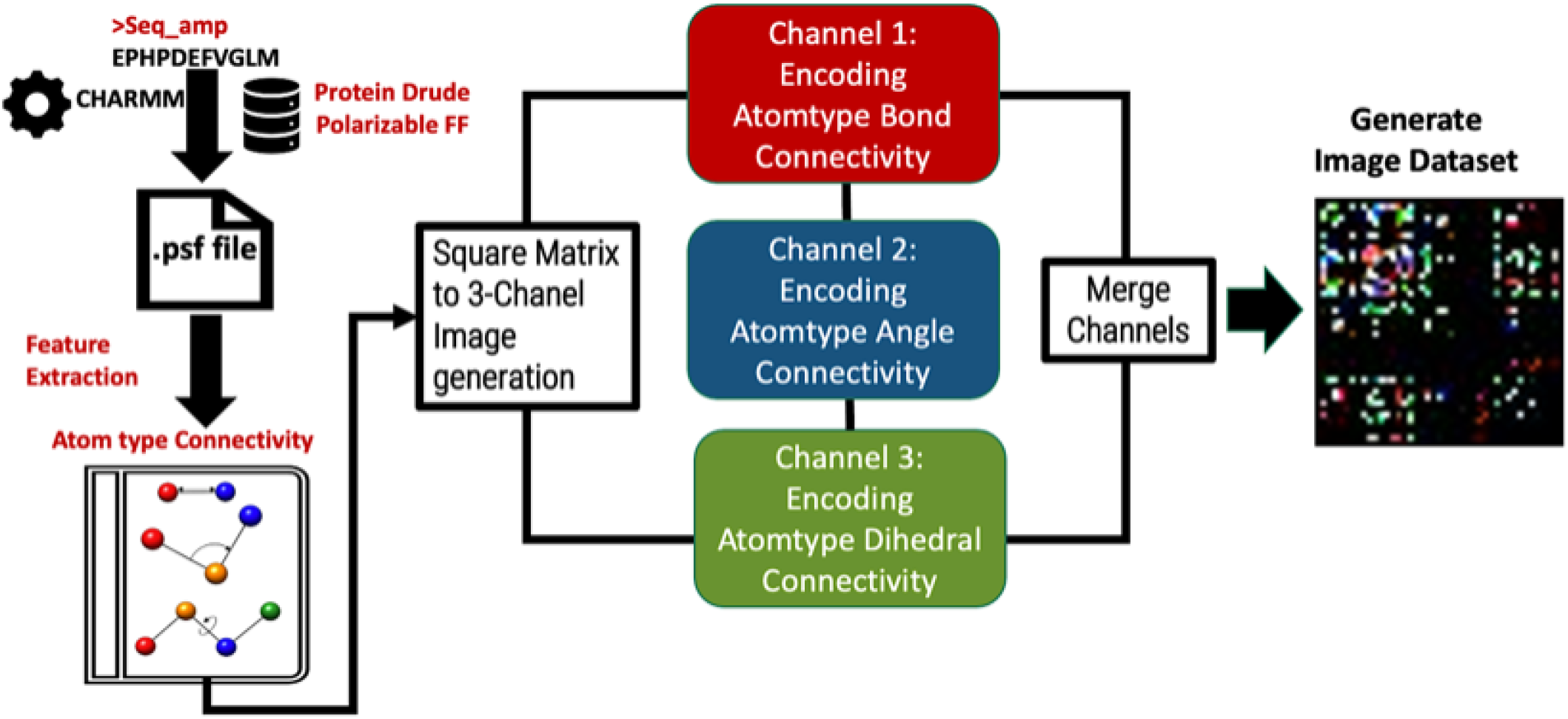
The framework of feature extraction and image generation for VGG16-AMP model.

## C. Training and Validation of VGG16-sAMP Model

VGG16[57] is a deep convolutional neural network that consists of five convolutional blocks with thirteen convolutional layers, five max-pooling layers, and three fully connected (FC) layers (Figure 2(a)). The first block consists of two convolutional layers with 64 feature kernel filters followed by max-pooling layer. The second block consists of two convolutional layers with 128 filters followed by max-pooling. The third block consists of three convolutional layers with 256 filters followed by maxpooling. The fourth and fifth block consist of 3 convolutional layers with 512 filters followed by max-pooling. Each of the convolutional layer is followed by RELU nonlinearity. Max-pooling is performed over a 2×2 pixel window with a stride of 2. The fifth block is followed by 3 fully connected (FC) dense layers. The output of the last pooling layer acts as input to the first FC layer. The first two FC layers consists of 4096 nodes and “ReLU” activation function and last FC layer consists of with 1000 nodes and SoftMax activation. The graphical representation of the VGG16 network is shown in Figure 3(a).

**FIG. 3:**
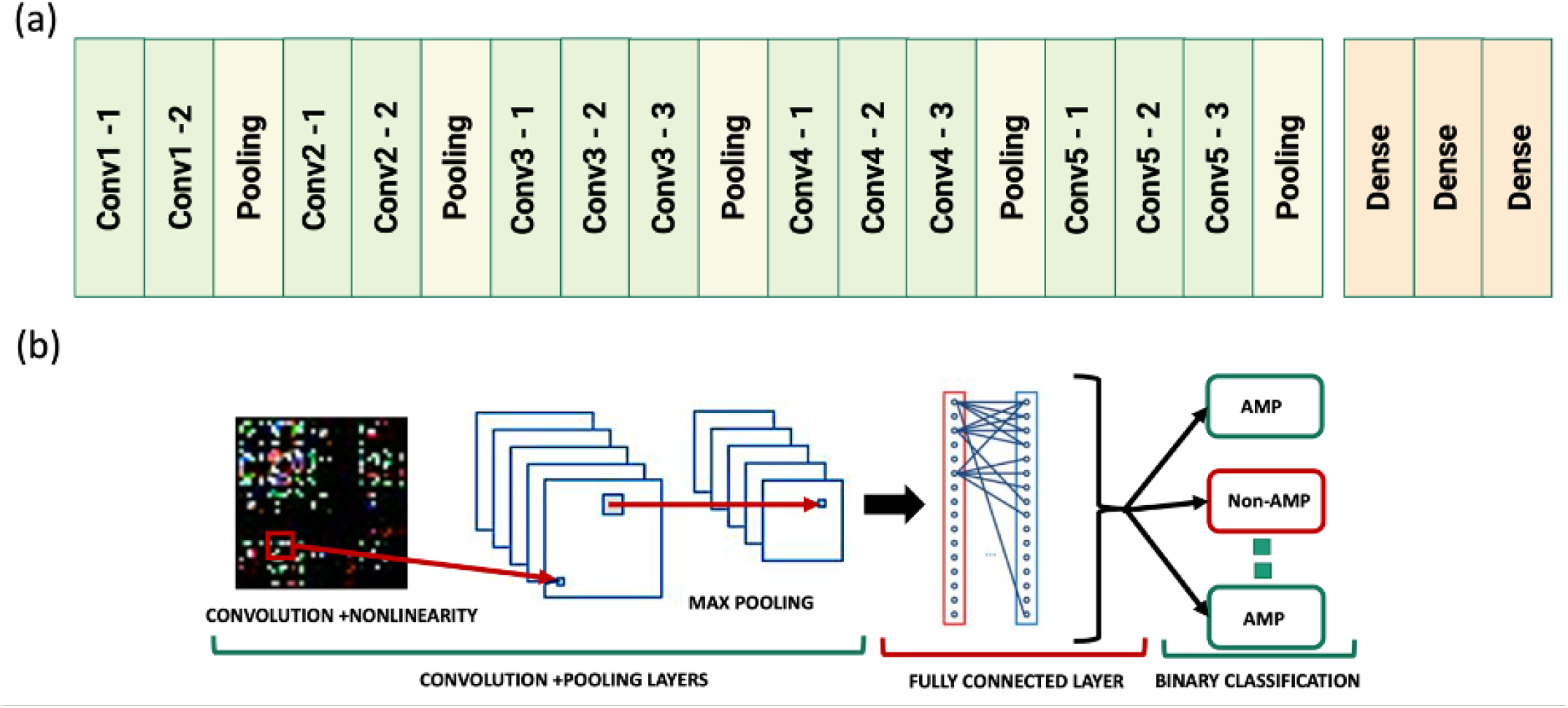
The graphical representation of (a) basic structure of VGG16 architecture, (b) prediction model of VGG16-AMP.

VGG16 model has been pretrained on the ImageNet[58] dataset. In this work, the fully connected layers of VGG16 are removed, and the following layers are added: (a) flatten layer, (b) FC dense layer with 4096 nodes and “RELU” activation function, (c) dropout layer with a rate of 0.7, (d) FC dense layer with 4096 nodes and “RELU” activation function, (e) dropout layer with a rate of 0.7, and (f) final FC dense layer with an output size of 1 and “sigmoid” activation function. Only the added fully connected layers has been trained to create the VGG16-AMP model. The graphical representation of prediction from VGG16-AMP is shown in Figure 3(b) and Figure S1 of SI File. The loss function used is mean square error and Adam optimizer is used to adjust model weights. The learning rate is set to 0.0001. The maximum number of epochs is set to 100, and the batch size is set to 24. A 10-times 10-fold cross-validation method was utilized to check the robustness of the VGG16-AMP network with chosen hyperparameters. Subsequently, the whole training dataset is used to develop the final VGG16-AMP model. The developed VGG16-AMP model is tested and compared with state-of-the-art-methods using benchmark dataset. The prediction architecture was implemented in Python using Keras[59] deep learning API and the TensorFlow[60] library.

## C. Evaluation of Performance

The prediction models are evaluated using seven performance indices including accuracy (ACC), sensitivity (Sn), specificity (Sp), Matthew’s correlation coefficient[61] (MCC), Cohen’s Kappa coefficient (Kappa), area under the Receiver Operating Characteristic (AUC-ROC) score, and area under the Precision-Recall (AUC-PR) score. The first five indices (ACC, Sn, Sp, and MCC) are defined as follows:

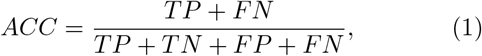

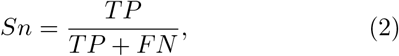

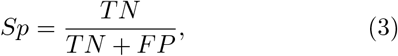

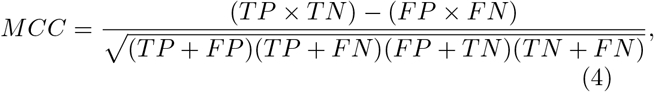

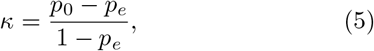

where,

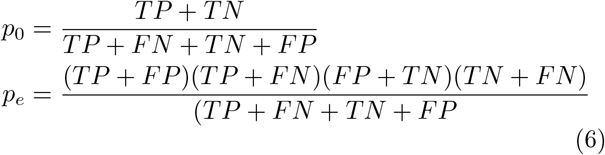

where TP, FP, TN, and FN denote the number of true positives, false positives, true negatives, and false negatives, respectively, and can be defined as:

[(a)]True Positive (TP): Both the target class and predicted class are positive. False Positive (FP): The target class is negative, but the predicted class is positive. True Negative (TN): Both the target class and predicted class are negative. False Negative (FN): The target class is positive, but the predicted class is negative.

Accuracy measures the fraction of correctly predicted classes[62]. Sensitivity measures the fraction of correct predictions in a positive class, while specificity measures the fraction of correct predictions in overall predictions of a negative class. MCC is a more reliable and comprehensive matrix[61, 63], as it considers all the four groups (TP, TN, FP, and FN) of the confusion matrix. The MCC matrix represents the correlation between the target and the predicted class values. MCC returns a value in the range of -1 (implies total disagreement between target and predicted class values) to 1(represents perfect prediction) and zero value infers random prediction. Cohen’s Kappa[64, 65] was first established to access agreement between two observers judging the same group of instances on a nominal scale with two or multiple classes. Similar to correlation coefficients, *κ* value may vary from -1 to +1, where zero indicates the degree of agreement anticipated by chance and one represents unanimous agreement between raters. The AUC-ROC[65] curve is a performance measure of classification problems at different thresholds. ROC is the probability curve and AUC represents the degree or degree of separation. It illustrates how the model can distinguish classes. The higher the AUC, the better the model for distinguishing the classes.

## III. RESULTS AND DISCUSSION

We aim to develop VGG16 based prediction model for AMPs using atomistic information from Drude polarizable FF. To do this, as a first step, we implemented 10fold cross-validation of binary classification with AMP-VGG16 10 times on a training dataset of 1,529 AMPs and 1,529 non-AMPs to demonstrate the robustness of the prediction model for external data. Figure 3 shows the comparative analysis of the prediction performance of VGG16 based prediction model with the existing prediction models trained on the same training dataset by 10 times 10-fold cross-validation. The 10-times 10-fold cross-validation results highlight that the VGG16-based prediction model outperformed the existing CNN, RF, or SVMlinear-based prediction models for the six performance indices (ACC, AUC-ROC, AUC-PR, Kappa, Sn, and MCC), as shown in Table S1 of SI file. However, the Sp value is higher for SVMlinear-based prediction models, followed by the VGG16-based prediction model. For the VGG16-based prediction model, the mean accuracy (ACC) is 0.8072 ± 0.01, the mean area under the ROC curve (AUC-ROC) is 0.8873 ± 0.01, the mean area under the precision-recall curve (AUC-PR) is 0.8750 ± 0.01, the mean Cohen’s Kappa coefficient (Kappa) is 0.6143 ± 0.02, the mean sensitivity (Sn) is 0.8747 ± 0.02, the mean specificity (Sp) is 0.7396 ± 0.01, and the mean Matthews correlation coefficient (MCC) is 0.6225 ± 0.02.

The chosen hyperparameters were then used to develop a prediction model using the whole training dataset. Figure 4 shows ROC curve and area under curve (AUC-ROC) for the training and benchmark dataset for VGG16-AMP model. For VGG16-AMP model, AUC-ROC values of the training and benchmark dataset are 0.94, and 0.86, respectively. For the proposed VGG16-AMP model, ACC, AUC-ROC, AUC-PR, Kappa, Sn, Sp, and MCC indices of the benchmark dataset are 0.809, 0.863, 0.863, 0.617, 0.798, 0.819 and 0.617, respectively. The proposed VGG16-AMP model, then compared with the existing state-of-the-art methods, including iAMP-2L24, iAMPpred27, AmPEP, AMP Scanner DNN30, RF-AmPEP3031, Deep-AmPEP3031 and LEV-KSVM[66] for benchmark dataset. The iAMP-2L prediction model was based on pseudo amino acid composition (PseAAC) and fuzzy K-nearest neighbor (FKNN) algorithm. The iAMPpred prediction model utilizes compositional, physicochemical, and structural features and SVM algorithm. The AmPEP prediction method was based on distribution patterns of amino acid properties and random forest algorithm. AMP Scanner DNN method was based on convolutional long-short-term memory (conv-LSTM) architecture, capturing position-invariant amino acid patterns along peptide sequences. RF-AmPEP30 and Deep-AmPEP30 models were developed using pseudo-K-tuple reduced amino acid composition (RAAC) with RF and convolutional neural network, respectively. The comparative analysis of the prediction performance of the proposed prediction models and above-mentioned state-of-the-art prediction models is shown in Figure 5. The proposed prediction model outperformed other classifiers for the benchmark dataset, indicating that the model has much greater discriminative power in terms of ACC, MCC, Kappa and Sp indices, as shown in Table S2 of SI file.

**FIG. 4:**
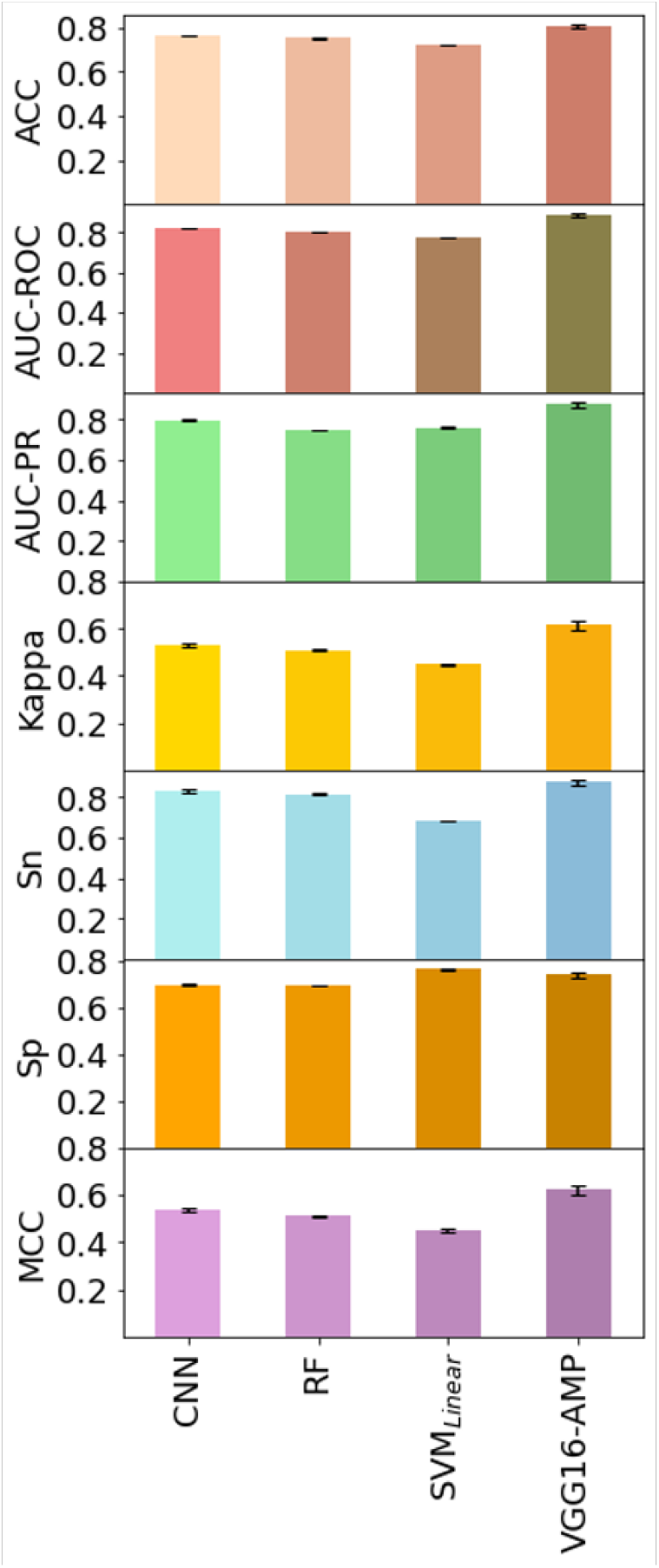
Comparison of VGG16-AMP-based prediction model with the existing prediction models trained on the same training dataset by 10 times 10-fold cross-validation.

**FIG. 5:**
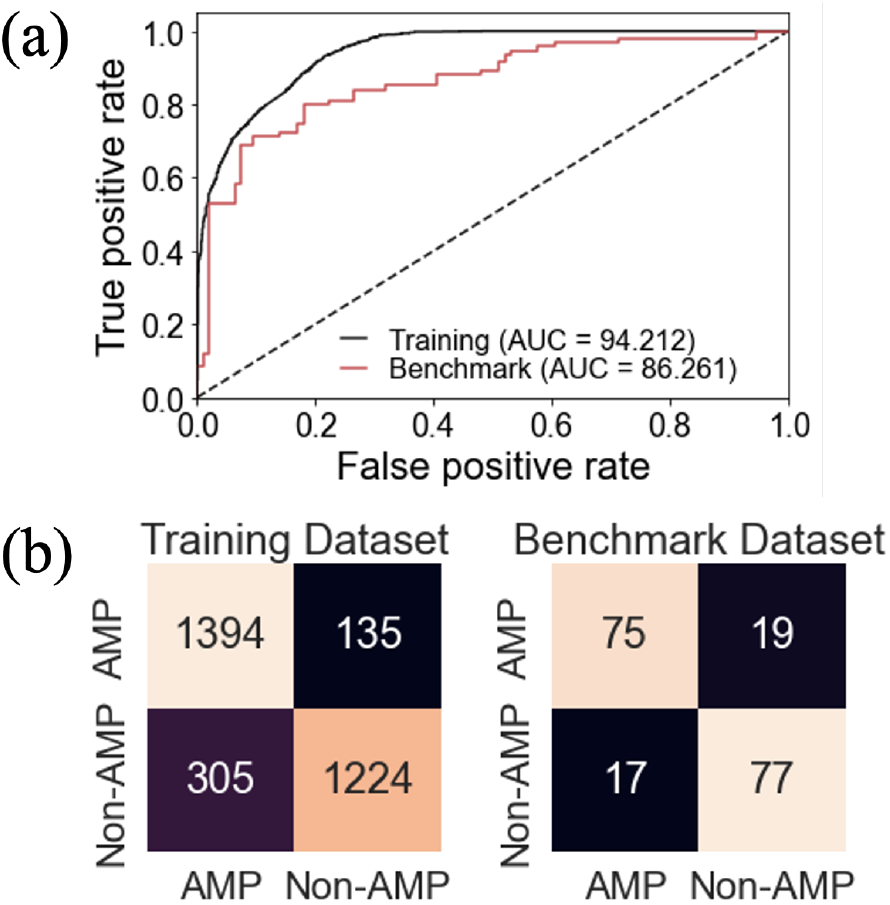
Figure showing (a) ROC curve, and (b) confusion matrices for training and benchmark dataset for VGG16-AMP model.

**FIG. 6:**
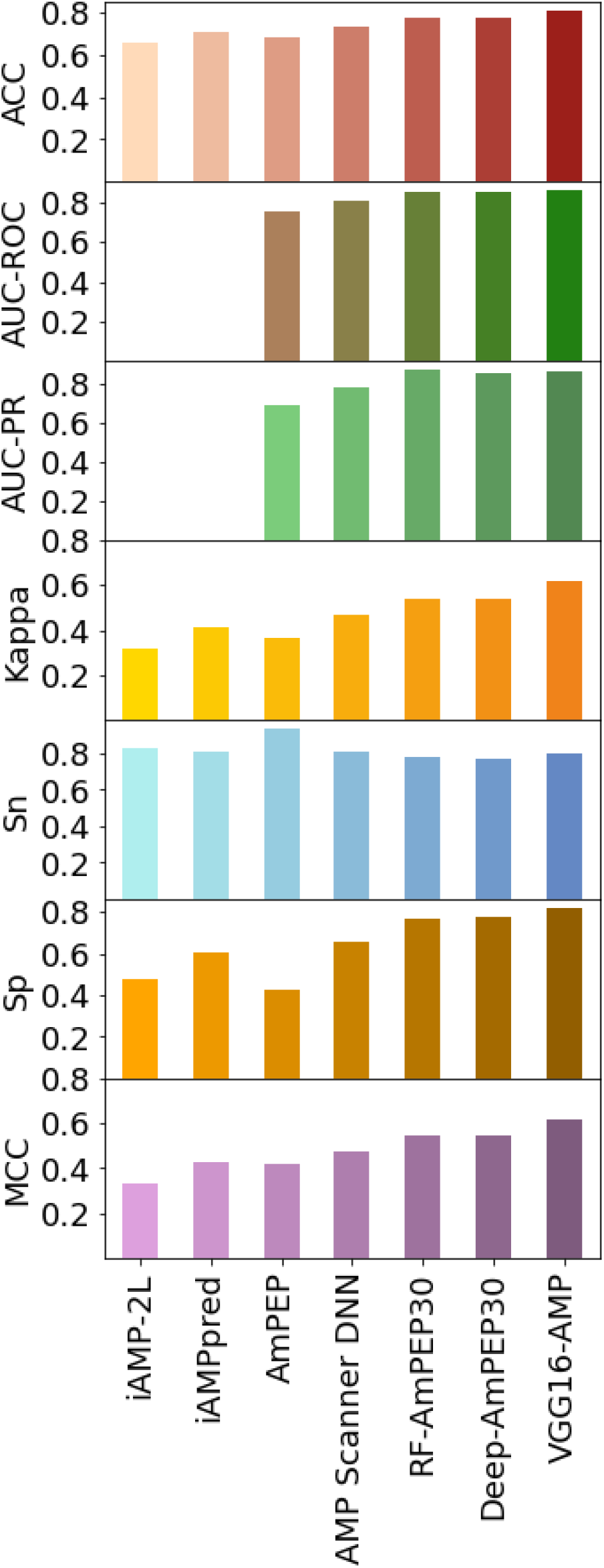
Comparison of VGG16-AMP-based prediction model with the existing state-of-the-art prediction models using the benchmark dataset.

## IV. CONCLUSION

This study successfully implemented transfer learning approach to the area of AMP prediction and recognition and proposed a new FF atom type-based feature encoding method to develop VGG16-AMP prediction model. In this work, we have assessed the efficiency of the VGG16 in conjunction with Drude atom type-based features as a model for sAMP prediction. The proposed VGG16-AMP classifier achieves better AMP recognition in comparison to existing state-of-the-art methods. To the best of our knowledge, this is the first instance where atomistic information has been used to predict small AMPs. By incorporating FF based atomistic information and VGG16 architecture, the proposed method automatically extracts expert-free features, hence eliminating the need of domain expert in feature encoding. Based on our results, we conclude that the proposed methodology yields reliable classifier of sAMPs. The present study highlighted that the transfer learning paradigm has great promise for protein classification and might have far-reaching implications for bioinformatics.

## V. KEY POINTS

- Proposed a novel computational protocol, sAMP-VGG16 for identifying short-antimicrobial peptides (sAMPs).
- The study is the first to utilize FF-based atomistic information to develop VGG16 based convolutional neural network for the protein classification problem.
- The FF-based atomistic features (bond, angle, and dihedral connectivities) can improve the model’s predictive performance for short-antimicrobial peptides (sAMPs).
- In Drude FF, an atom can adopt a distinct atomtype definition premised on its connectivity along with the formal charge state. So, the atom-type annotation differentiates atoms depending on their chemical environment in the peptide residues, providing effective features for prediction model development.
- The sAMP-VGG16 achieved high accuracy and outperformed several existing state-of-the-art methods for predicting sAMPs.

## VI. DATA AVAILABILITY

Our code, models, and curated datasets are publicly available at this repository https://doi.org/10.6084/m9.figshare.23123429.v1

## VII. COMPETING INTERESTS

The authors declare that the study was done in the absence of any commercial ties that might be perceived as a possible conflict of interest.

## VIII. AUTHOR CONTRIBUTIONS STATEMENT

PP and AS conceived the study. PP developed the prediction model and analyzed the results. PP prepared the manuscript. AS supervised the study and edited the manuscript. The final version of the article has been reviewed and approved by all authors.

## XI. ACKNOWLEDGMENTS

The authors thank the anonymous reviewers for their valuable suggestions. PP acknowledges the Department of Biotechnology (DBT), Government of India for the DBT-Research Associate Fellowship (Award Letter No. DBT/2022/January/06).

## Supporting Information File

**Figure S1.**
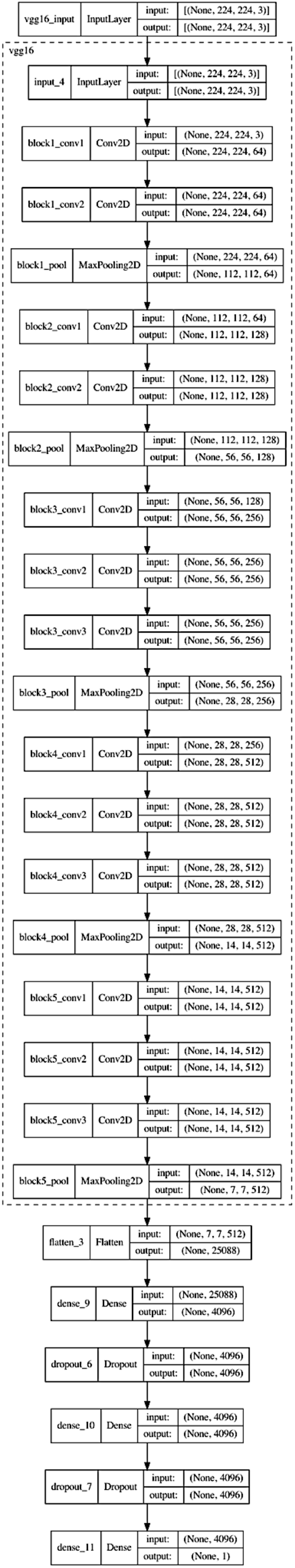
The graphical summary of VGG16-AMP model.

**Table S1.** Comparison of VGG16-based prediction model with existing prediction models by 10 Times 10-fold cross-validation

| Methods | ACC | AUC-ROC | AUC-PR | Kappa | Sn | Sp | MCC | Reference |
| --- | --- | --- | --- | --- | --- | --- | --- | --- |
| CNN | 76.50 ± 0.37 | 82.48 ± 0.20 | 79.55 ± 0.5 | 53.00 ± 0.74 | 83.35 ± 0.86 | 69.65 ± 0.67 | 53.51 ± 0.78 | Yan et.al. |
| RF | 75.42 ± 0.23 | 80.58 ± 0.09 | 74.85 ± 0.21 | 50.84 ± 0.46 | 81.55 ± 0.42 | 69.28 ± 0.22 | 51.23 ± 0.48 | Yan et.al. |
| SVM<sub>linear</sub> | 72.37 ± 0.24 | 77.90 ± 0.07 | 76.08 ± 0.14 | 44.75 ± 0.49 | 68.36 ± 0.29 | 76.39 ± 0.42 | 44.95 ± 0.50 | Yan et.al. |
| VGG16-AMP | 80.72 ± 0.97 | 88.73 ± 1.10 | 87.50 ± 1.28 | 61.43 ± 1.95 | 87.47 ± 1.56 | 73.96 ± 1.34 | 62.25 ± 2.01 | This study |

**Table S2.**
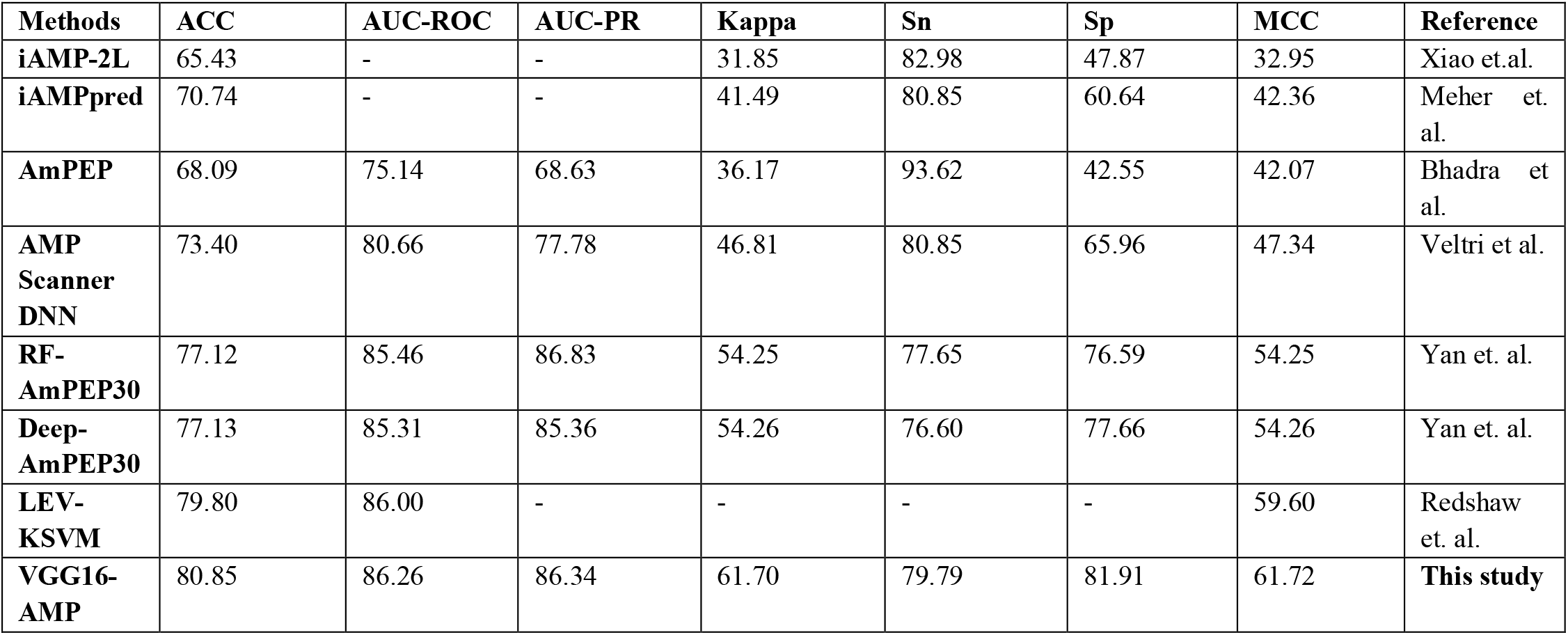
Comparison of VGG16-based prediction model with existing prediction models using the Benchmark Dataset.

## Notes

https://doi.org/10.6084/m9.figshare.23123429.v1

### Competing Interest Statement

The authors have declared no competing interest.

